# Sex Differences in Acute Responses to Psychedelics: Evidence for Greater Subjective Intensity and Impairment in Female Participants

**DOI:** 10.64898/2026.07.08.737179

**Authors:** Natasha L Mason, Eline CHM Haijen-Bongers, Kim PC Kuypers, Aline Frick, Stefan W Toennes, Pablo Mallaroni, Johannes G Ramaekers

## Abstract

Serotonergic psychedelics are advancing as psychiatric treatments, yet acute responses vary between individuals and the contribution of sex, a fundamental biological variable, remains largely unexamined. We pooled two double-blind, placebo-controlled randomized studies in healthy volunteers (*N* = 94 drug observations; 43 male, 51 female) comparing psilocybin 15 mg, 2C-B 20 mg, and LSD 50 µg. Linear mixed models tested sex differences in acute subjective effects (visual analogue scales), retrospective altered-states ratings (5D- and 11-ASC), empathy (Multifaceted Empathy Test), and peak (Cmax) and total (AUC) drug exposure, with treatment and sex as fixed factors and age as a covariate. Female participants reported numerically higher subjective ratings than male participants on most measures. After adjustment for age, sex differences were significant on five measures, most robustly for feeling under the drug’s influence, reduced vigilance, and impaired control and cognition, with medium-to-large effects. These effects were largely consistent across the three drugs. No sex differences emerged on any empathy measure or in peak drug concentrations, indicating that female’s stronger responses occurred at equivalent drug exposure. Female participants may experience more intense acute subjective effects and greater perceived impairment under psychedelics, independent of age and not explained by drug exposure. These preliminary findings, implicating pharmacodynamic rather than pharmacokinetic mechanisms, have implications for dosing, informed consent, and safety monitoring, and underscore the need to treat sex as a biological variable in adequately powered psychedelic trials.

**Trial register information:** Study 1, Sub(acute) Neuropsychopharmacological Profiling of 2C-B vs Psilocybin, https://onderzoekmetmensen.nl/en/trial/50024, NL-OMON50024.

Study 2 The effect of LSD on neural synchrony, prosocial behavior, and relationship quality, https://onderzoekmetmensen.nl/nl/trial/53689, NL-OMON53689.

## Introduction

Serotonergic psychedelics such as psilocybin and lysergic acid diethylamide (LSD) are advancing through clinical trials for multiple psychiatric disorders (1), yet acute responses vary markedly between individuals (2, 3), and the biological sources of this variability remain poorly understood — with consequences for efficacy (4–9), tolerability, and safety (10–12). One fundamental but largely overlooked biological variable is sex (13). Females experience adverse drug reactions roughly twice as often as males, in part because fixed dosing yields higher drug exposure in females (14). Moreover, psychedelics produce their effects through agonism at the serotonin 2A (5-HT2A) receptor (15). Ovarian hormones modulate the serotonergic system, including 5-HT2A receptor density (16–18), and fluctuate across the menstrual cycle. Sex-related biology could thus shape both the subjective intensity of psychedelics and associated socio-cognitive effects such as empathy.

A growing preclinical literature reports sex-dependent psychedelic effects, including greater psilocybin-induced head-twitch responses — a 5-HT2A–mediated behavioral marker of psychedelic potential — in female than male mice (19, 20), alongside sex-differentiated effects on reward, affective, developmental, and social behaviour (21–24); yet sex differences in acute psychedelic response have rarely been examined directly in humans, despite the routine inclusion of female participants in psychedelic trials (13, 25).

Despite the routine inclusion of female participants in psychedelic trials, sex differences in acute psychedelic response have rarely been examined in both clinical and preclinical psychedelic research (13, 25, 26). This gap is relevant because emerging preclinical work suggests that sex influences serotonergic-mediated behavioral sensitivity to psychedelics (26). Whether a similar sex difference is evident in humans remains unclear, with implications for dosing, informed consent, and safety monitoring (13). To address this gap, we pooled data from two placebo-controlled experimental studies spanning three psychedelics to test for sex differences in acute subjective drug effects and empathy.

## Methods

A detailed description of the experimental procedure is provided in the Supplementary Methods and briefly summarized here.

### Study design

This pooled analysis combined two double-blind, placebo-controlled studies in healthy volunteers with prior psychedelic experience. Study 1 used a randomized crossover design (2C-B 20 mg, psilocybin 15 mg, placebo); Study 2 used a randomized within-subject design (LSD 50 µg, placebo). Procedures are detailed in the original reports and the Supplementary Methods. Analyses focused on drug-induced changes for psilocybin, 2C-B, and LSD; depending on the measure they were derived from the active session directly or by subtracting the placebo session. Both studies were approved by the local Medical Ethics Committee and conducted in accordance with the Declaration of Helsinki (27), and took place at Maastricht University. Participants were enrolled between between 2021 and 2022 for Study 1, and between 2022 and 2026 for Study 2.The present data were part of larger clinical trials, both registered on onderzoekmetmensen (Study 1: https://onderzoekmetmensen.nl/en/trial/50024, NL-OMON50024, 06-05-2021; Study 2: https://onderzoekmetmensen.nl/nl/trial/53689, NL-OMON53689, 13-09-2022).

### Participants

The pooled sample comprised 72 participants (32 male, 40 female): Study 1 included 22 (11 female) and Study 2 included 50 (29 female). Because 22 participants contributed two active sessions (psilocybin and 2C-B), the dataset comprised 94 active-drug observations (43 male, 51 female).

### Outcome Measures

Acute subjective effects were rated on visual analogue scales (feeling high, the drug’s influence, good drug effect, bad drug effect, drug liking) and summarized as Emax per item, accommodating the drugs’ differing time courses. Retrospective ratings used the 5- and 11-dimensional Altered States of Consciousness scales (5D-ASC, 11-ASC (31, 32)), which are explicitly referenced to the normal waking state and therefore index drug-induced change directly (32). Empathy was measured with the Multifaceted Empathy Test (MET; cognitive and emotional empathy for positive and negative stimuli) (33). As the MET is not baseline-referenced, it was expressed as placebo-corrected change (active minus placebo) to isolate the drug effect. Plasma drug concentrations (PK) were sampled at predefined time points throughout the testing days. Because the compounds exhibited differing concentration–time profiles, peak concentration (Cmax) was used as a time-independent exposure metric to enable comparison across drugs. Additionally, AUC was calculated per participant using the linear trapezoidal rule over a common 0–300 min post-administration window, harmonized across the three studies to maximize the number of participants with complete profiles. As absolute exposure differs by orders of magnitude across psilocin, 2C-B, and LSD, both Cmax and AUC were standardized (z-scored) within drug before pooling, so that the sex contrast reflects relative exposure on a common scale rather than absolute concentration.

### Statistics

Statistical analysis was conducted in IBM SPSS Statistics 28. Outcome variables of the 5D- & 11D ASC, VAS, MET and PK were analyzed using a full factorial linear mixed model (LMM). Fixed effects included Treatment (psilocybin, LSD, or 2C-B), Sex (female or male), their interaction (Treatment * Sex), and age (mean-centred), with a random intercept for participants to account for the non-independence of repeated observations. Age was included as a covariate because it differed between male and female participants, to adjust the sex contrast for age; each model was estimated both with and without this covariate, and unadjusted and age-adjusted estimates are reported (Table S1), with the age-adjusted models treated as primary. As previous publications point to valence-specific effects of psychedelics on empathy (34), this analysis was repeated for negative and positive stimuli on the MET separately. Demographic information (age, education, lifetime use of psychedelics, BMI) was compared between males and females via independent-samples t-tests.

Statistical significance was set at a P value of less than .05. For the sex contrast, Cohen’s *d* was the female-minus-male estimated marginal mean difference divided by the model-implied total SD, the square root of the summed residual and random-intercept variances; 0.2/0.5/0.8 denote small/medium/large (35, 36). Given the exploratory aim, no multiple-comparison correction was applied and emphasis is placed on effect sizes. Findings are considered preliminary and intended to inform future confirmatory research.

## Results

### Demographics

Male participants were significantly older than female participants, t(70) = 2.59, p = .012; males: M = 26.97, SD = 5.39; females: M = 24.05, SD = 4.15, but the groups did not differ in BMI (p=.325), lifetime psychedelic use (p=.148) or education (p=.295). Because age differed by sex and younger individuals may experience stronger acute effects (37), age was included as a covariate.

### Female participants reported stronger acute subjective drug effects

Female participants scored numerically than males on every item (Figure 1). After adjusting for age, the sex difference was significant for Feeling Under the Drug’s Influence (F_(1, 54.91)_ = 6.63, p = .013, d = 0.68) and Good Drug Effect (F_(1, 55.73)_ = 4.10, p = .048, d = 0.50), and non-significant for Feeling High (F_(1, 57.28)_ = 2.95, p = .091, d = 0.45), Drug Liking (F_(1, 38.99)_ = 0.70, p = .408, d = 0.20), and Bad Drug Effect (F_(1, 41.20)_ = 0.25, p = .620, d = 0.13). A Treatment × Sex interaction emerged for Feeling Under the Drug’s Influence (F_(2, 42.49)_ = 3.33, p = .045), reflecting a female–male difference that was most pronounced under psilocybin (Supplementary Results); no other interaction was significant (all p > .05). Age was not associated with any VAS outcome (all p ≥ .226). See Table S1 for unadjusted and age-adjusted estimates.

**Figure 1.**
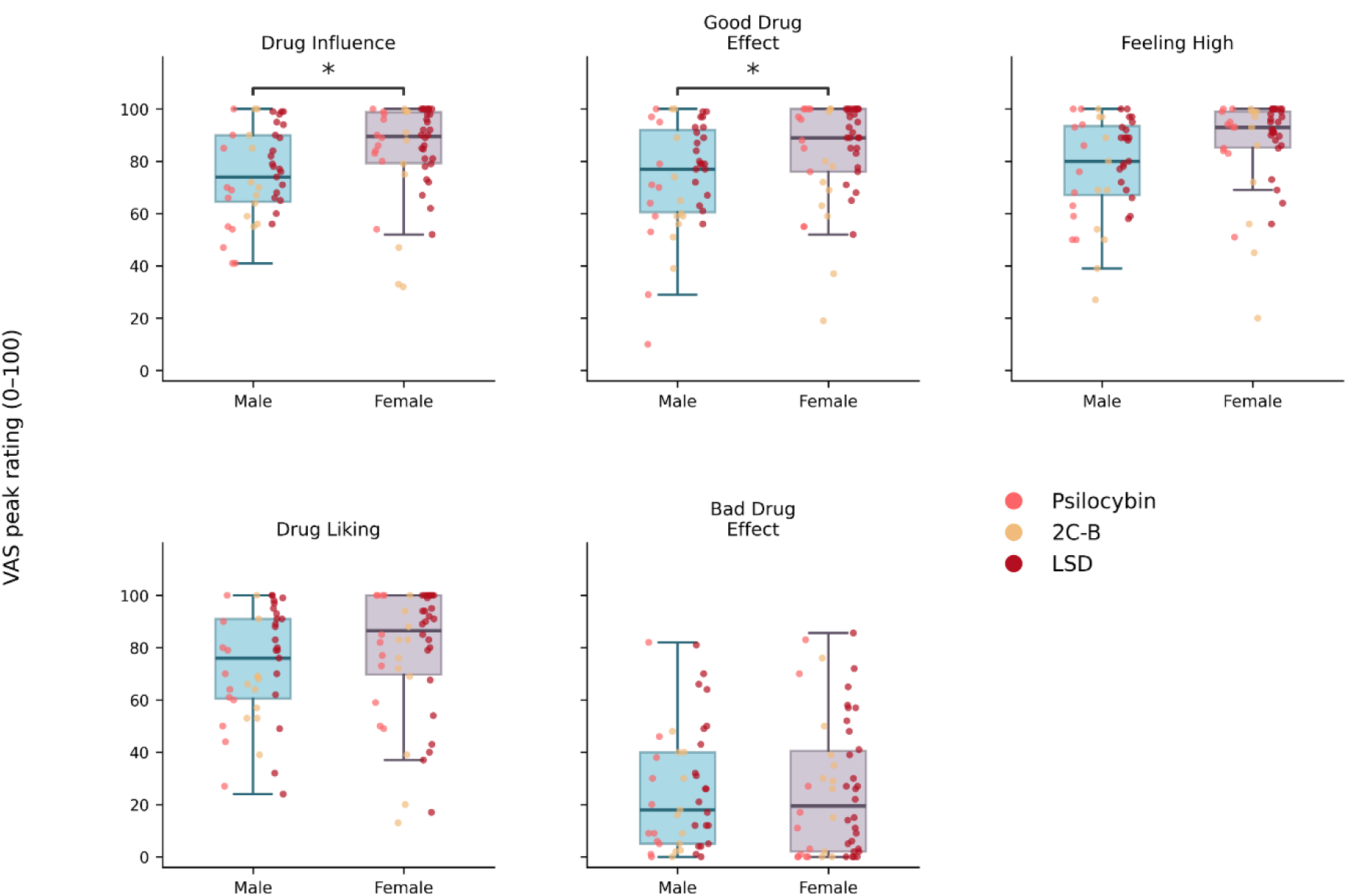
Sex differences in acute subjective drug effects. Peak drug effects (Emax) on five visual analogue scales, by sex. Boxes show the median and interquartile range (IQR); whiskers extend to the furthest data points within 1.5×IQR of the quartiles; points are coloured by drug (psilocybin, 2C-B, LSD). Asterisks denote a significant age-adjusted sex difference (* p < .05, ** p < .01)..

### Females reported stronger retrospective ratings of drug experience

Female participants scored numerically higher than males on all five 5D-ASC dimensions (Figure 2; Figure S1a for spider plot). After adjusting for age, the sex difference was significant for Reduction of Vigilance (F_(1, 37.85)_ = 8.15, p = .007, d = 0.73) and Anxious Ego Dissolution (F_(1, 42.17)_ = 4.16, p = .048, d = 0.51), and non-significant for Visionary Restructuralization (F_(1, 49.42)_ = 1.18, p = .282, d = 0.31), Oceanic Boundlessness (F_(1, 43.44)_ = 1.15, p = .289, d = 0.29), and Auditory Alterations (F_(1, 44.76)_ = 0.03, p = .865, d = 0.04). A Treatment × Sex interaction emerged for Reduction of Vigilance (F_(2, 28.94)_ = 3.47, p = .045), with the female–male difference significant under psilocybin and 2C-B but not LSD (Supplementary Results); no other interaction was significant (all p > .05). Age was independently associated with four of the five dimensions (all p ≤ .041) but not Reduction of Vigilance (p = .088). See Table S1 for unadjusted and age-adjusted estimates.

**Figure 2.**
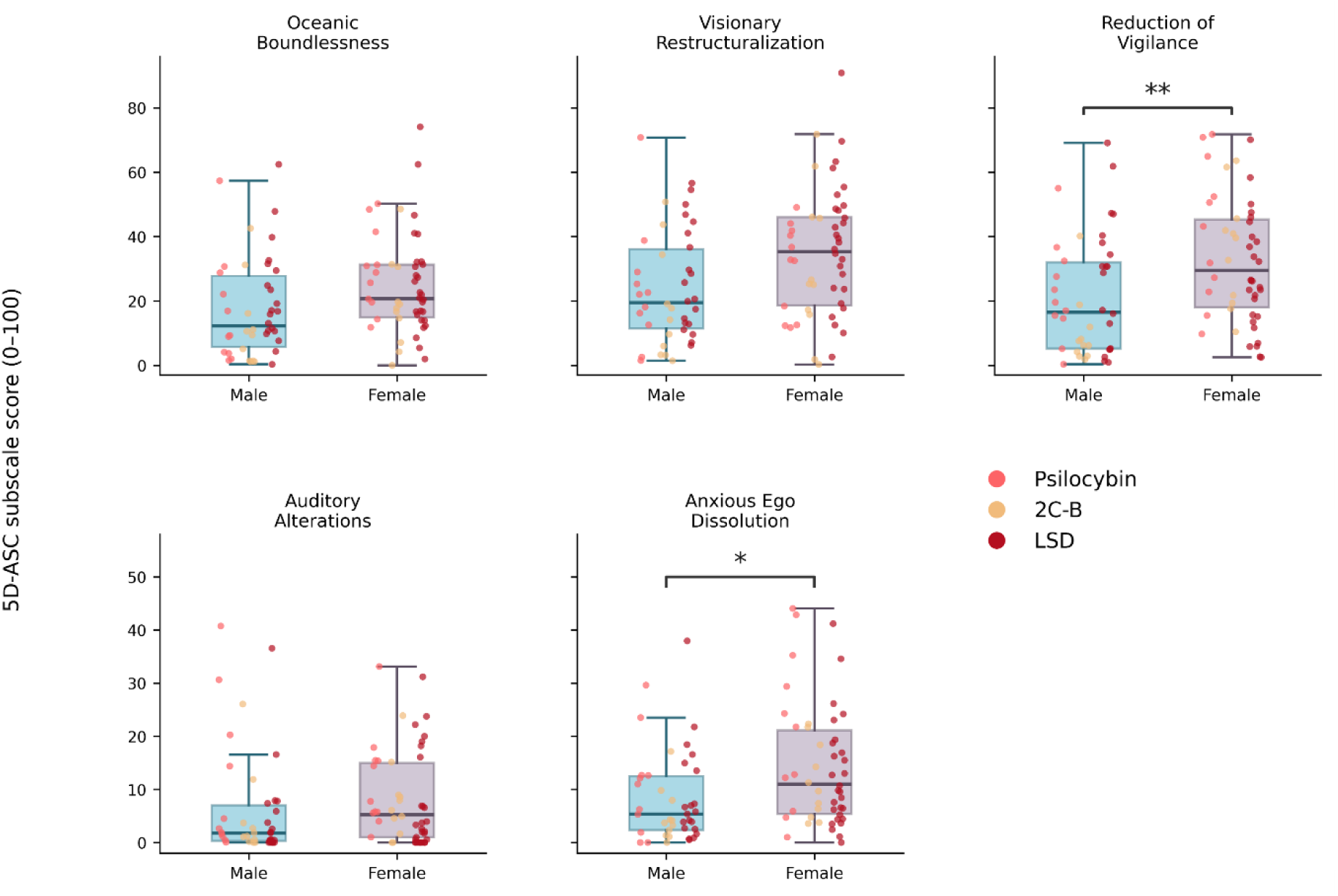
Sex differences in retrospective ratings of the 5D-ASC. Subscale scores by sex for the five 5D-ASC dimensions. Boxes show the median and IQR; whiskers extend to 1.5×IQR; whiskers extend to the furthest data points within 1.5×IQR of the quartiles; points are coloured by drug (placebo excluded). Asterisks denote significant age-adjusted sex differences (* p < .05, ** p < .01).

On the 11-ASC, female participants scored numerically higher than male participants on most factors (Figure 3; Figure S1b for spider plot). After adjusting for age, this difference was significant for Impaired Control and Cognition, F_(1, 27.35)_ = 5.83, p = .023, d = 0.59. No other sex main effect reached significance (all p ≥ .098). Age was independently associated with six of the eleven 11-ASC dimensions (Experience of unity, Impaired control and cognition, elementary imagery, insightfulness, complex imagery, blissful state; all p ≤ .032). See Table S1 for unadjusted and age-adjusted estimates.

**Figure 3.**
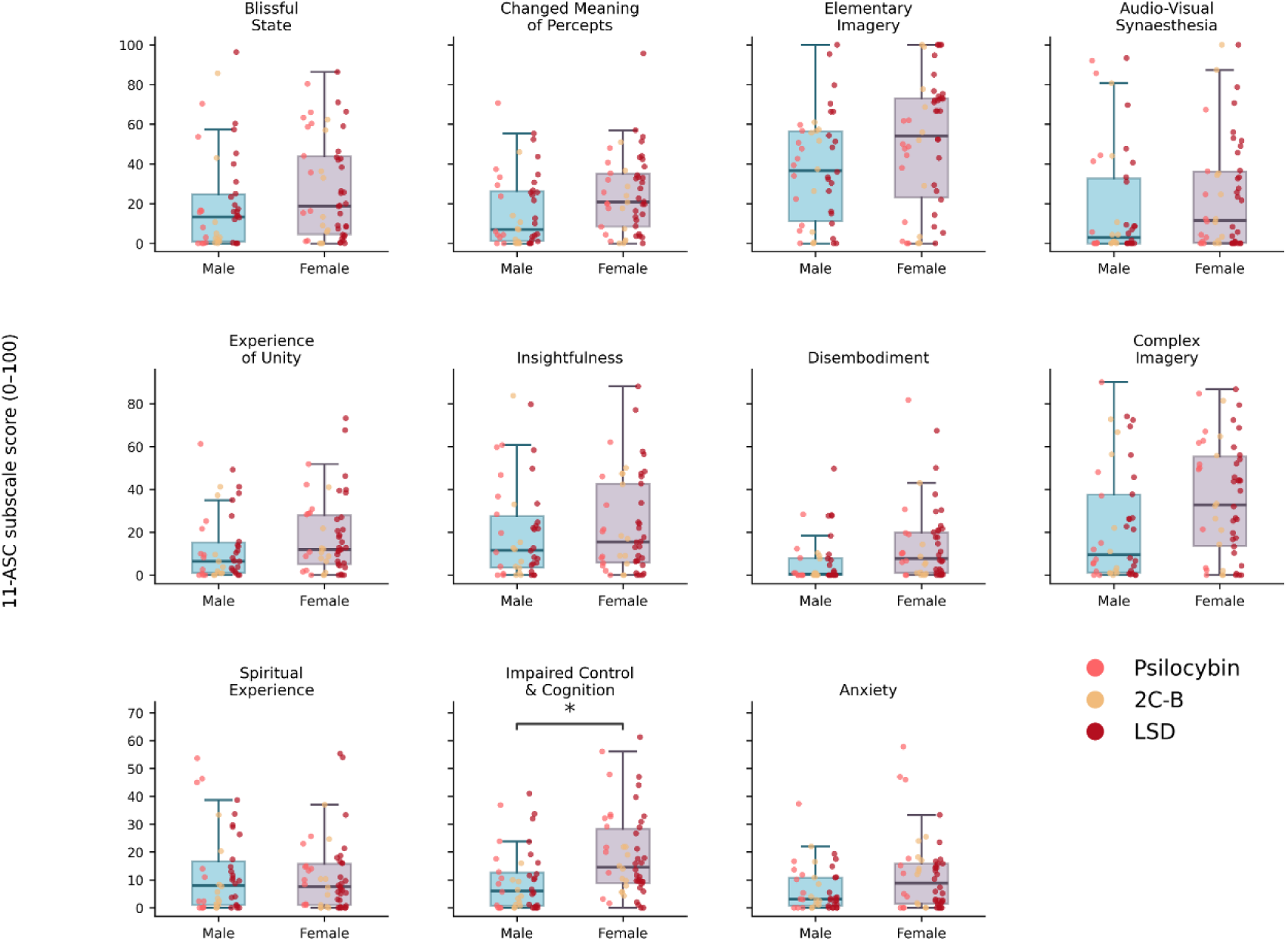
Sex differences in retrospective ratings of the 11-ASC. Subscale scores by sex for the eleven 11-ASC factors. Boxes show the median and IQR; whiskers extend to 1.5×IQR; whiskers extend to the furthest data points within 1.5×IQR of the quartiles; points are coloured by drug (placebo excluded). Asterisks denote significant age-adjusted sex differences (* p < .05, ** p < .01)..

### Acute effects of psychedelics on empathy are not sex-dependent

On the Multifaceted Empathy Test, there were no significant sex differences in cognitive empathy, implicit emotional empathy, or explicit emotional empathy, whether stimuli were pooled or analyzed separately by valence (all p ≥ .208). Age was not associated with any empathy measure (all p ≥ .147). See Table S1 for full statistical tests and confidence intervals.

### Greater subjective intensity in females was not explained by peak drug or total exposure

Peak plasma concentrations (Cmax) and AUC of the active compounds, psilocin, 2C-B, and LSD, were standardized within drug and compared by sex. After adjusting for age, neither Cmax, F_(1, 46.92)_ = 0.39, p = .533, nor AUC, F_(1, 35.97)_ = 0.73, p=0.398 differed between sexes. The Treatment × Sex interaction was non-significant for Cmax F_(2, 30.40)_ = 2.28, p = .120 and AUC F_(2, 26.46)_ = 1.13, p=.337. Age was not associated with Cmax (p = .159) or AUC (p=.481). Results were unchanged without age adjustment: Cmax, F_(1, 45.52)_ = 0.03, p = .873, AUC, F_(1,35.76)_ = 0.44, p=.510.

## Discussion

The present study investigated sex differences in acute subjective responses and empathy measures following administration of psychedelics, using pooled data from two experimental psychedelic studies spanning three psychedelics. Overall, female participants reported numerically higher subjective ratings across most measures, with the largest difference from male participants on feeling under the influence of the drug, reduced vigilance, and impaired control and cognition; smaller but significant sex differences also emerged for good drug effect and anxious ego dissolution. No sex differences emerged for any empathy measure. Sex effects were largely consistent across substances, the two exceptions: the female-male difference in feeling under the drug’s influence was most pronounced under psilocybin, and the difference in reduced vigilance was present under psilocybin and 2C-B but not LSD. Although age accounted for substantial variance in acute responses, consistent with previous findings (2), these sex effects remained significant after adjustment for age. Critically, these differences did not appear to be driven by drug exposure: neither peak plasma concentrations nor total drug exposure differed by sex, indicating that female participants reported more intense effects at comparable drug levels. Together, these results suggest that female subjects may experience more pronounced acute subjective drug effects, including greater perceived impairment, under a psychedelic.

These outcomes are clinically relevant. Reduced vigilance, impaired control and cognition, and anxious ego dissolution may indicate that female subjects experienced the acute psychedelic state as more disruptive to alertness, cognitive clarity, and perceived control. Such experiences may be therapeutic and meaningful when adequately supported but may also affect tolerability and may increase the need for reassurance and the level of psychological support required during dosing sessions (38, 39). This pattern converges with earlier controlled 3,4-methylenedioxymethamphetamine (MDMA) trials. Liechti and colleagues (40) reported that female participants experienced more intense subjective MDMA effects than male participants, particularly perceptual changes, thought disturbance, and fear of loss of body control, as well as more frequent acute adverse effects and sequelae. Although MDMA is pharmacologically distinct from classic psychedelics such as psilocybin and LSD, the overlap in phenomenological domains is notable.

The most straightforward explanation for the greater acute intensity and impairment reported by female subjects would be pharmacokinetic. Because psychedelic trials use fixed rather than weight-adjusted doses, female subjects may reach higher peak plasma concentrations than male subjects; across drug classes, such female-biased pharmacokinetics parallel the roughly two-fold higher rate of adverse drug reactions in female subjects, a relationship that is not explained by body weight (14). Our data, however, do not support this account: neither peak plasma concentrations nor total exposure (AUC) differed significantly by sex. The greater subjective intensity reported by female participants thus occurred at equivalent peak and cumulative exposure, arguing against a purely pharmacokinetic mechanism. This inference is bounded by our measure: Cmax and AUC capture peak and total exposure but not unbound exposure, and we did not assess metabolism or plasma protein binding (41), which could still differ by sex. Nonetheless, female participants responded more strongly despite comparable drug levels. These findings align with the largest predictor analysis of the LSD experience to date, in which total LSD exposure (plasma AUC) was associated with CYP2D6 genotype and dose, but not sex (42). The absence of a sex difference in both total exposure (previous study) and peak concentration (current study) suggests that the sex effect on intensity sits downstream of drug exposure. Notably, that study did not identify sex as a significant predictor of the subjective effects of LSD. That said, sex was one of nineteen predictors tested against twenty-nine outcomes with correction across all 551 tests, a configuration in which a medium-sized, focal sex effect such as the one found in this study would be readily masked among many demographic variables.

Attention therefore shifts to pharmacodynamic mechanisms-differences in responsiveness at the drug’s target rather than in exposure. Classic psychedelics produce their subjective effects through agonism at the serotonin 2A (5-HT2A) receptor (15), and ovarian hormones act directly on this system: estradiol increases 5-HT2A receptor density in cortical and limbic regions in rodents (16) and in human prefrontal cortex (43), alongside broader modulation of serotonin synthesis, transport, and receptor expression (13). Greater 5-HT2A availability could therefore amplify the subjective response to a given level of drug exposure. Consistent with this, preclinical studies report stronger LSD and psilocybin-induced head-twitch responses in female than male mice across studies (12, 19, 44–46). Psychedelic responses appear to vary across ovarian hormones. Tylš and colleagues (47) found that psilocin-induced behavioral effects are blunted during high estrogen phases (estrus and proestrus) whereas Zylko and colleagues (19) reported stronger head twitches in response during diestrus and proestrus. Additionally, progesterone has been shown to reduce LSD-induced effects, and female rodents showed blunted responses to a 5-HT1a agonist during high-estrogen phases of the estrous cycle (48–50). These findings suggest that sex and ovarian hormones may not increase or decrease psychedelic sensitivity globally but rather shape specific behavioral and physiological response domains.

Because ovarian hormones like estradiol and progesterone fluctuate across the menstrual cycle, psychedelic responsivity may also vary across cycle phases, with shifting estradiol up- or down-regulating sensitivity to 5-HT2A-mediated effects. Low-ovarian-hormone phases such as the periovulatory and perimenstrual phases can additionally coincide with heightened stress sensitivity and negative affect that may shape the psychological “set” in which the experience unfolds (51). These hormonal accounts predict that hormonal status contributes to within-sex variability, reinforcing the need to assess menstrual-cycle phase in future work.

These observations echo a broader lesson from psychopharmacology: sex differences in drug response are common but often recognized late. Female patients tend to respond better to SSRIs and male patients to tricyclics — an effect concentrated in premenopausal patients, implicating the same serotonergic–hormonal systems considered here (52). And where such differences have been characterized rather than averaged over, they have changed practice: zolpidem’s recommended dose was halved for female patients after they were found to reach higher plasma concentrations and greater next-day impairment (53), and policies now require sex to be treated as a biological variable in research design and analysis (54). Psychedelic trials, still early in establishing their design conventions, are unusually well placed to incorporate these lessons prospectively rather than retrospectively.

For psychedelics specifically, these differences also carry direct clinical weight. Because the intensity of the acute altered state predicts therapeutic improvement following psychedelics (55), sex differences in its magnitude may shape both tolerability and efficacy. This complicates dosing: although greater perceived impairment in female participants might argue for lower or individualized doses, reducing the dose risks blunting the experiential features that mediate benefit. Therefore, dose optimization should be guided by clinical outcome rather than acute intensity alone. More immediately, informed consent and pre-session preparation should convey that female participants may experience more intense effects, including greater impairment and potentially anxiety; acute monitoring and psychological support should be resourced accordingly; and trials should be powered to test sex as a biological variable, report sex-disaggregated outcomes, and document hormonal status, including menstrual-cycle phase, hormonal contraceptive use, and reproductive stage.

This study represents an initial step toward understanding sex differences in psychedelic responses, but several limitations apply. It is a secondary analysis of pooled data from two studies, neither designed to assess sex differences, and substantial within-sex variability among female participants remains unexplained — likely in part owing to menstrual-cycle phase, which was not assessed. Hormonal status was not analysed here and represents priorities for direct testing of the mechanisms proposed above. The sample also consisted of healthy volunteers with prior psychedelic experience, which may limit generalizability to clinical populations. This limitation may be particularly relevant for female patients, as patient samples may differ in menstrual cycle regularity, hormonal contraceptive use, reproductive stage, and stress- and symptom-linked hormonal dynamics (56–58). Sex-related differences observed in healthy participants may therefore be amplified, attenuated, or qualitatively different in patient samples. Despite these limitations, the results demonstrate the value of pooled datasets for investigating sex differences in psychedelic effects.

In conclusion, this pooled analysis found that female participants reported stronger acute subjective drug effects, including greater perceived impairment, whereas no sex differences emerged in empathy. These findings are preliminary but point to meaningful sex-related variability in subjective psychedelic responses that warrants investigation in adequately powered, sex-sensitive studies.

## Acknowledgements

Study 1 was supported by the Dutch Research Council (Grant No. 406.18. GO.019 [to JGR]). Study 2 was supported by the Mind Science Foundation (2023) Brainstorm award (to NLM, PM, JGR). NLM is financially supported by the Dutch Research Council (NWO, grant number VI.Veni.231G.011). Thank you to all participants for the time, effort and trust, to Cees van Leeuwen for medical supervision, and to all interns for their help in participant recruitment, screening, and data collection. During the preparation of this work, the author(s) used Claude (Anthropic) to refine, edit, and format human-written text; to format and generate presentation-ready tables of research data (e.g., assembling results tables from statistical output); and to produce code used to generate figures. All AI-assisted output was reviewed, verified, and edited by the author(s), who take full responsibility for the content of the publication.

## Author Contributions

Conceptualization: NLM, EHB, KPCK. Data acquisition: NLM, PM. Analysis: NLM, SWT. Writing-Original draft: NLM, EHB, AF. Writing-Review & Editing: all authors. Supervision: NLM. Funding Acquisition: NLM & JGR.

## Data availability

Data can be made available upon request to N.L.M. and J.G.R. and a data use agreement executed with Maastricht University.

## Supplementary Information for

**Table S1.**
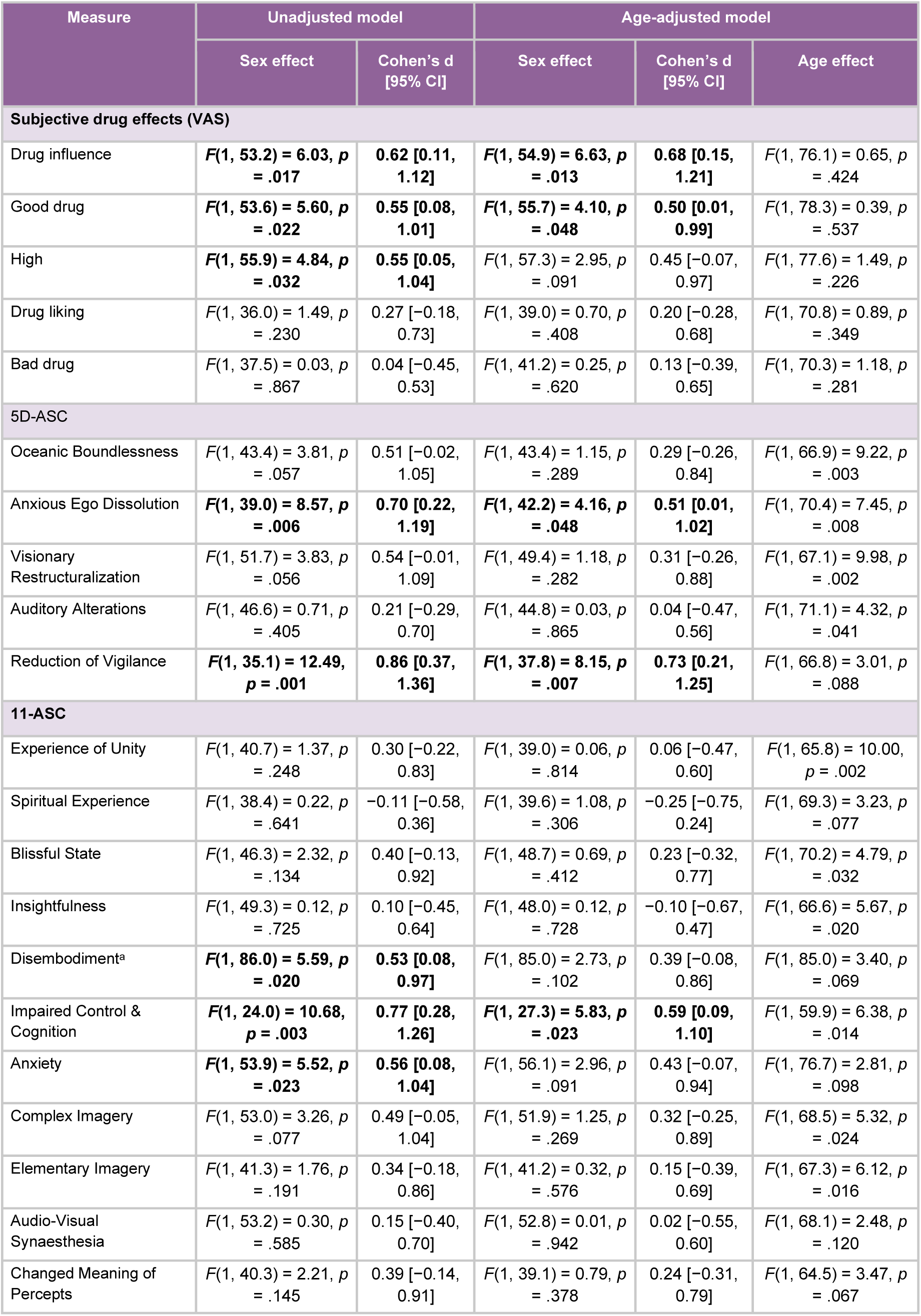
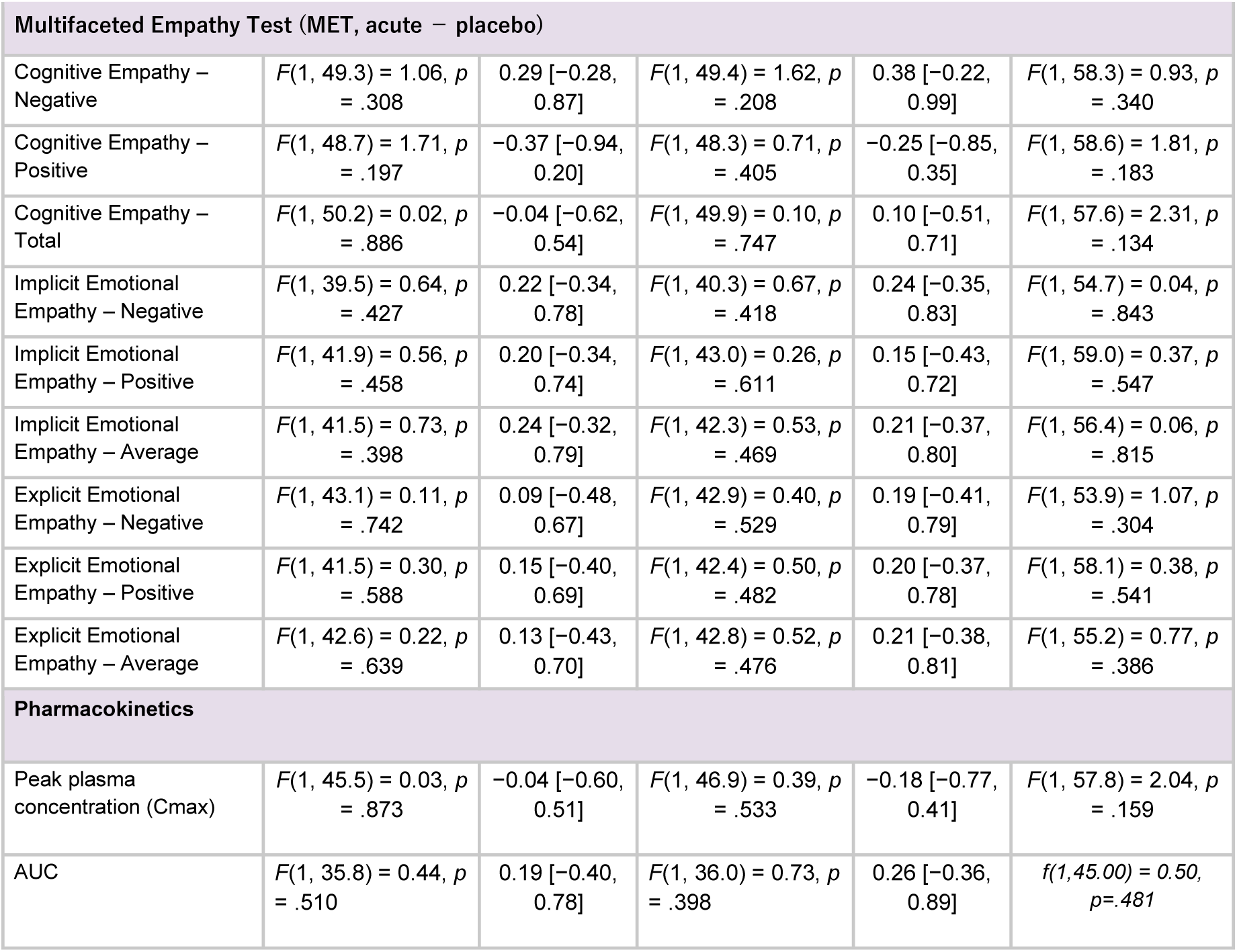
Sex differences before and after adjustment for age, across subjective drug effects (VAS), 5D-ASC, 11-ASC, and Multifaceted Empathy Test (MET) outcomes.

**Figure S1.**
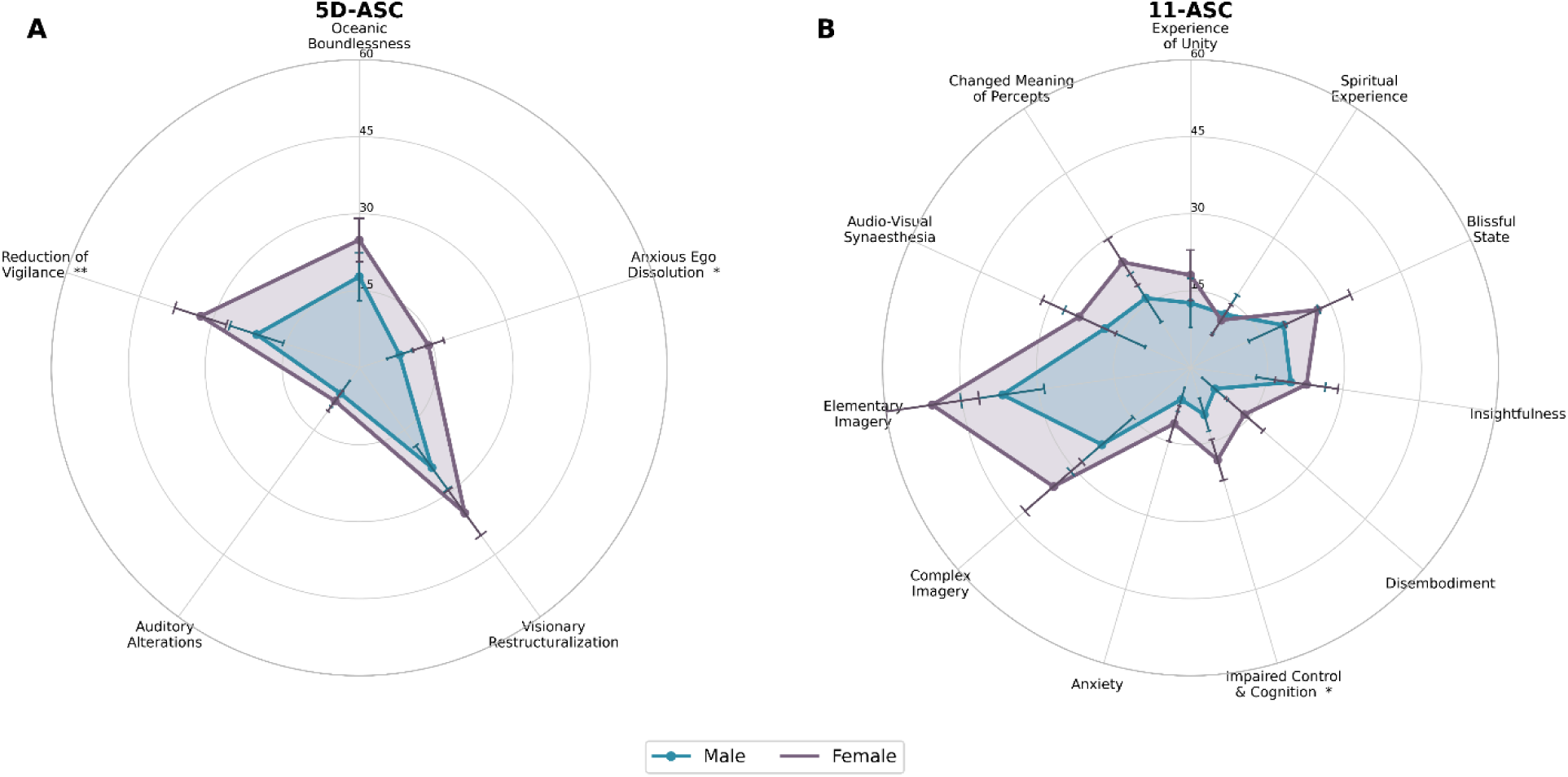
Altered-states consciousness profiles by sex. Mean (± 95% CI) subscale scores for males and females across **(A)** the 5D-ASC and **(B)** the 11-ASC, pooled across active drugs (placebo excluded). Asterisks mark significant age-adjusted sex differences from the linear mixed models (* p < .05, ** p < .01).

**Figure S2.**
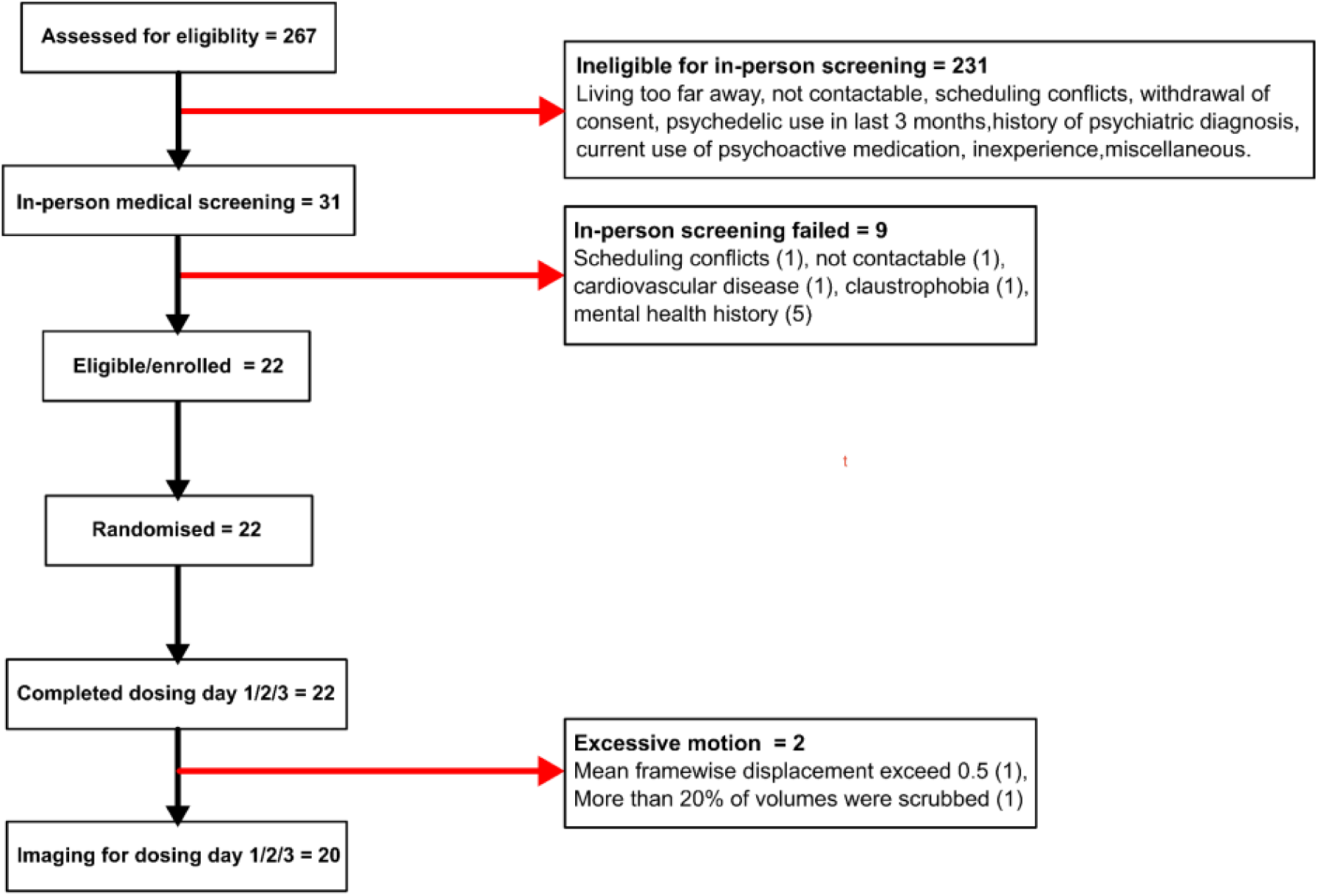
CONSORT flow chart for Study 1. As previously published in prior publications.

**Figure S3.**
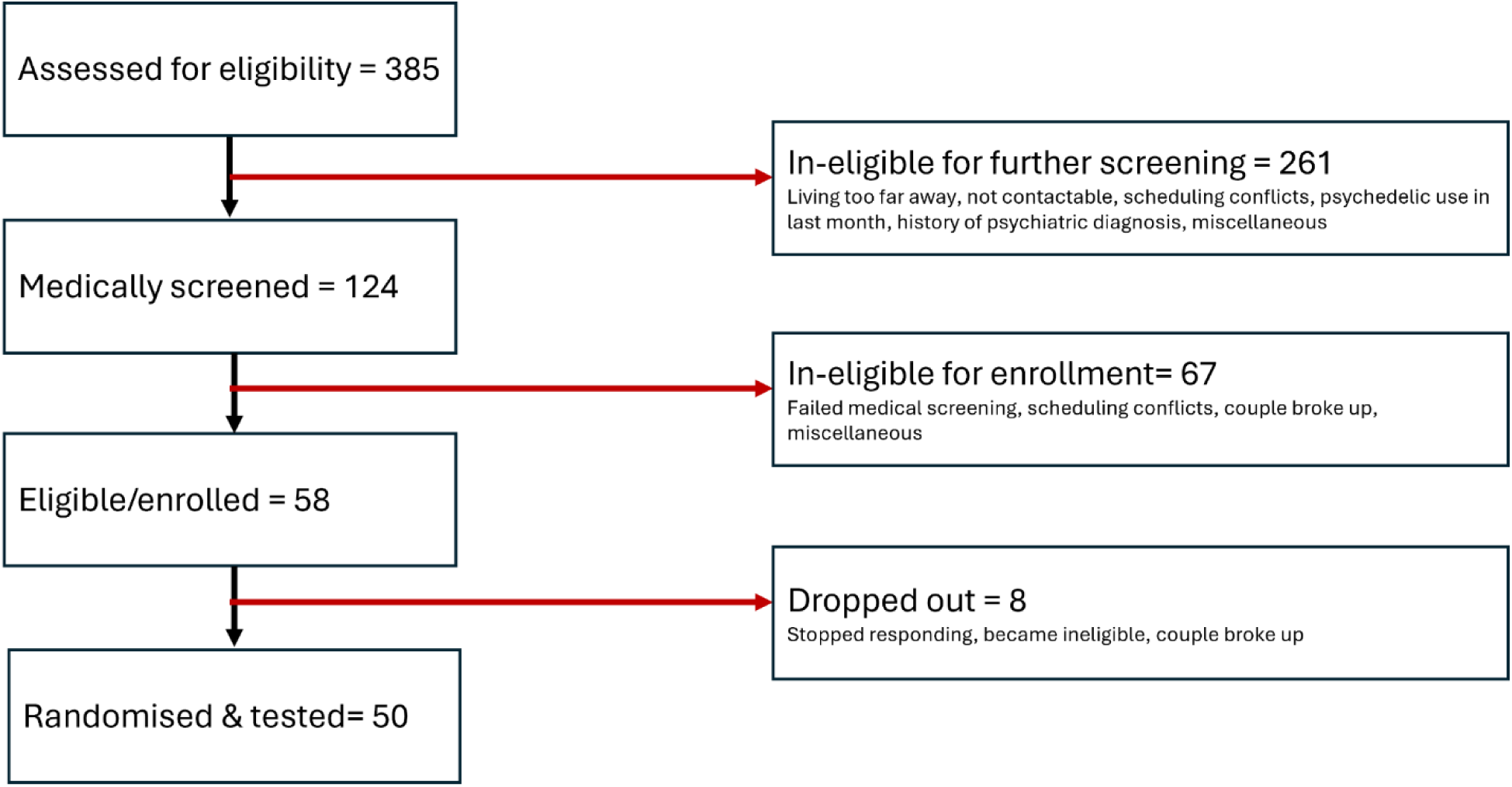
CONSORT flow chart for Study 2.

### Supplementary Methods

#### Ethics and approvals

Both studies were conducted in accordance with the Declaration of Helsinki (1964, amended in Fortaleza, Brazil, October 2013) (1) and the Dutch Medical Research Involving Human Subjects Act (WMO), and were approved by the Academic Hospital and University’s Medical Ethics Committee. A permit for obtaining, storing, and administering 2C-B, LSD, and psilocybin was obtained from the Dutch Drug Enforcement Administration. All participants were fully informed of the procedures, possible adverse reactions, legal rights and responsibilities, expected benefits, and their right to voluntary termination without consequences.

#### Recruitment and consent

In both studies, participants were recruited through online advertisements and underwent an initial online pre-screening to assess preliminary eligibility, including prior substance use, general health status, and availability. Eligible participants were invited to an online briefing, during which study procedures, potential risks, legal rights and responsibilities, and the right to withdraw without consequence were explained. Participants then provided written informed consent before completing structured inventories assessing psychiatric history and drug use.

#### Eligibility

Criteria were largely harmonized across the two studies. In both studies, participants were healthy adults between the ages of 18-40 years old, with previous psychedelic experience but no psychedelic use within the past 3 months, BMI between 18 and 28 kg/m², no major medical or psychiatric disorder, no history of substance abuse or dependence, no pregnancy or lactation, no previous serious adverse reactions to psychedelics, and reliable contraception for female participants. Study-specific differences were that Study 1 (psilocybin vs placebo) excluded MRI contraindications, whereas Study 2 (LSD) required participants to be in a steady relationship for at least 6 months.

Before inclusion, participants underwent an in-person medical screening by an independent study physician or psychiatrist. This included a review of physical and mental health, vital signs, body weight, resting electrocardiogram, physical examination, urine toxicology, urinalysis, hematology, serum chemistry, and liver function tests. Female participants completed a pregnancy test. Participants who met all criteria were enrolled and remunerated for participation.

#### Randomization and blinding

An experimenter who was not responsible for treatment randomization or preparation, recruited all participants. A separate experimenter, who did not come in direct contact with the subjects, allocated treatment following Latin-square counterbalancing. This latter experimenter was also responsible for preparing the treatment (psilocybin, 2CB, LSD, placebos), and giving the treatment to the data collector in a masked cup (psilocybin, 2CB) or vial (LSD). The masked cup/vial ensured that neither the data collector nor the participant would be unblinded as to what treatment the participant was receiving.

#### Experimental sessions

Each experimental session lasted approximately 7 hours. Participants were instructed to eat a light breakfast before arrival and to abstain from psychedelics for at least 3 months, alcohol for at least 24 hours, and other drugs of abuse for at least 7 days before each testing period. Participants were also asked to avoid caffeine and nicotine on test days. On arrival, recent drug and alcohol use were assessed using urine toxicology and a breathalyzer, and female participants completed a pregnancy test. Testing proceeded only if all results were negative. During the session, participants remained under supervision. Participants were discharged only once the experimental team judged them fit to leave.

#### Drug administration

For Study 1, synthetic (powder) formulations of 2C-B and psilocybin were employed. 2C-B was obtained from Duchefa Farma B.V., Haarlem, the Netherlands. Psilocybin was obtained from THC Pharm GmbH, Frankfurt, Germany. Participants were informed prior to the study they were to receive 20 mg 2C-B, 15 mg psilocybin, or placebo on 3 separate occasions. Each intervention was administered orally, in a closed cup either containing 200 mL of the vehicle bitter lemon (a bittering agent, placebo) or bitter lemon and psilocybin/2C-B (powder) to mask any potential taste. Blinding was assessed retrospectively at the end of each dosing day.

For Study 2, LSD was obtained from Apotheke Dr. Hysek AG. Participants were told they would receive LSD (50 μg) and placebo on 2 separate occasions. LSD (50 μg) was administered in a small amount of alcohol (ethanol, 0.6mL/96% Vol.). Placebo was 0.6 mL of ethanol in an identical vial. The content of the vial was put in a syringe, which was used to deliver directly into the participant’s mouth.

#### Sample size calculation

This study is a pooled secondary analysis of data collected across 2 previously conducted placebo-controlled, within-subjects trials. As such, no a priori power analysis was performed for the present sex-difference comparison; the sample size was instead determined by the total number of participants available across the pooled datasets.

#### Outcome Measures

##### Acute subjective effects (VAS)

Throughout each test day, participants rated feeling high, feeling the drug’s influence, good drug effect, bad drug effect, and drug liking on visual analogue scales. Because the three drugs differ in temporal profile, each item was summarized per participant as Emax — the maximum post-drug rating— capturing peak acute response independent of the timing of peak effects.

##### Retrospective ratings of the drug experience

At the end of the test day, when drug effects had subsided, participants completed the 5-Dimensional Altered States of Consciousness Rating Scale and its 11-factor version. The 5D-ASC comprises oceanic boundlessness, anxious ego dissolution, visionary restructuralization, auditory alterations, and reduction of vigilance; the 11-ASC comprises experience of unity, spiritual experience, blissful state, insightfulness, disembodiment, impaired control and cognition, anxiety, complex imagery, elementary imagery, audio-visual synaesthesia, and changed meaning of percepts. Because the scale assesses changes relative to a person’s normal waking state, no additional baseline correction was applied.

##### Multifaceted Empathy Test (MET)

The MET comprises 40 photographs of people in emotional states (50% positive, 50% negative). Cognitive empathy was indexed by the number of correctly selected emotion words (from four options per image). Implicit (indirect) emotional empathy was indexed by ratings (1–9) of how aroused each image made the participant feel, and explicit (direct) emotional empathy by ratings (1–9) of how concerned they felt for the depicted person. Each index was computed separately for positive and negative stimuli and averaged across valence. For each active condition, placebo-corrected change scores were calculated by subtracting the placebo-day score from the active-day score. The MET has shown good-to-satisfactory validity and reliability (2) and sensitivity to psychedelics (3).

##### Pharmacokinetics

Blood samples were collected at predefined time points before and after administration to quantify plasma drug concentrations.

For Study 1, analysis of 2cb was via serum (200 µl), extracted with 1 ml of ethyl acetate/methyl tertiary-butyl ether (80:20, v/v) after addition of 0.2 ml phosphate buffer pH 9 and 50 µl of internal standard (acetonitrile containing 0.5 ng of 2C-B-d6 from Toronto Research Chemicals, Toronto, Canada). Analysis of psilocin in serum was performed according to (59). Serum (200 µl) was extracted with 1 ml of ethyl acetate after addition of phosphate buffer pH 9, 20 ng psilocin-d 10 and 10 µl of 0.1 M ascorbic acid for stabilization (4).

For both analysis streams, the organic phase was evaporated and reconstituted with 100 µl of 0.1 % formic acid/acetonitrile (80:20, v/v). The analysis of (5 µl 2C-B, 2 µl psilocin) was performed on an Agilent (Waldbronn, Germany) LC-MS/MS system consisting of a 1290 Infinity Liquid Chromatograph coupled via JetStream Electrospray Interface (ESI) to a G6460A Triple Quadrupole Mass Spectrometer. Analytes were separated on a Kinetex® 2.6 µm XB-C18 100 Å LC column (100 x 2.1 mm) plus corresponding guard column from Phenomenex (Aschaffenburg, Germany) at 30 °C. Gradient elution at a flow rate of 0.5 ml/min using 0.01% formic acid containing 5 mM ammonium formate (A) and acetonitrile containing 0.1 % formic acid (B) started with 5 % B, increased to 95 % B during 4 min and was held for 2 min. Source parameters were: gas temperature 300 °C, gas flow 11 l/min, nebulizer 45 psi, sheath gas temperature 400 °C, sheath gas flow 12 l/min and capillary voltage 3500 V. Detection was performed in the multiple reaction monitoring mode (m/z, collision energy in parentheses, quantifier underlined): 2C-B-d6: 266→249.0 (8V); 2C-B 260→243 (8V); 228 (20V); psilocin-d10: 215→66 (12), psilocine 205→58 (12); 205→160 (16). Seven calibration standards were prepared in the range 0.2 – 50 ng 2C-B and five calibration standards in the range 1 – 100 ng/ml for psilocin per ml human serum and were analysed with the samples. Calibrations were linear (regression coefficients >0.99).

For Study 2, analysis of LSD was via serum (500 µl), extracted with 2.5 ml of 1-chlorobutane/diethyl ether (1:1, v/v) after addition of 0.5 ml 1M phosphate buffer pH 9.5 and 50 µl of internal standard (acetonitrile containing 2.5 ng of LSD-d3 from Cerilliant, Round Rock, Texas, US). The organic phase was evaporated and reconstituted with 100 µl of 0.1 % formic acid/acetonitrile (80:20, v/v). The analysis of 5 µl was performed on an Agilent (Waldbronn, Germany) LC-MS/MS system consisting of a 1290 Infinity II Liquid Chromatograph coupled via JetStream Electrospray Interface (ESI) to a G6495D Triple Quadrupole Mass Spectrometer. Analytes were separated on a Kinetex® 2.6 µm XB-C18 100 Å LC column (100 x 2.1 mm) plus corresponding guard column from Phenomenex (Aschaffenburg, Germany) at 50 °C. Gradient elution at a flow rate of 0.5 ml/min using 0.01% formic acid containing 5 mM ammonium formate (A) and acetonitrile containing 0.1 % formic acid (B) started with 20 % B, increased to 50 % B during 1.5 min and further to 100 % B during 2.5 min. Source parameters were: gas temperature 250 °C, gas flow 13 l/min, nebulizer 20 psi, sheath gas temperature 400 °C, sheath gas flow 12 l/min and capillary voltage 4000 V. Detection was performed in the multiple reaction monitoring mode (m/z, collision energy in parentheses, quantifier underlined): LSD-d3: 327.2→226.1 (28V); LSD 324.2→208.1 (32V); 223.1 (28V). Six calibration standards were prepared in the range 5 – 2500 pg per ml human serum and were analysed with the samples. Calibrations were linear (regression coefficients >0.99).

### Supplementary Results

Decomposition of the significant Treatment × Sex interaction for feeling under the drug’s influence revealed that the sex difference was driven by psilocybin: females scored 23.1 points higher than males (F_(1, 83.27)_ = 10.43, p = .002), with no significant sex difference under 2C-B (Δ = 3.4, F_(1, 83.27)_ = 0.21, p = .636) or LSD (Δ = 7.5, F_(1, 83.32)_ = 2.34, p = .124).

A significant Treatment × Sex interaction was also observed for Reduction of Vigilance. Here the sex difference was present under psilocybin (Δ = 16.5, F_(1, 81.95)_ = 4.55, p = .036) and 2C-B (Δ = 22.9, F_(1, 81.95)_ = 8.76, p = .004), but not under LSD (Δ = −0.4, F(1, 82.08) = 0.01, p = .936).

A significant main effect of Treatment was observed for good drug effect, F_(2, 50.99)_ = 5.93, p = .005, feeling high, F_(2, 46.75)_ = 4.42, p = .017, and drug liking, F_(2, 35.84)_ = 3.53, p = .040. Bonferroni-corrected pairwise comparisons indicated that ratings were higher following LSD than 2C-B for good drug effect, p = .003, feeling high, p = .013, and drug liking, p = .035. No other drug comparisons reached statistical significance.

A significant main effect of Treatment was observed for Anxious Ego Dissolution, F_(2, 35.42)_ = 4.49, p = .018, and Oceanic Boundlessness, F_(2, 28.24)_ = 3.50, p = .044. Bonferroni-corrected pairwise comparisons indicated that Anxious Ego Dissolution scores were higher following psilocybin than 2C-B, p = .020; for Oceanic Boundlessness, no pairwise comparison reached significance. On the 11-ASC, significant main effects of Treatment were observed for Impaired Control and Cognition, F_(2, 22.74)_ = 4.01, p = .032, and Anxiety, F_(2, 48.92)_ = 3.43, p = .040. Bonferroni-corrected pairwise comparisons indicated that Impaired Control and Cognition scores were higher following psilocybin than 2C-B, p = .040, and that Anxiety scores were higher following psilocybin than LSD, p = .035. No other main effect of Treatment reached significance (all p ≥ .057).

There were no significant main effects of Treatment on any empathy measure, all p ≥ .183.

